# Deciphering the dynamics of Cyanobacteria-Phage in a natural lake: Insights from a decade-long investigation

**DOI:** 10.1101/2025.02.19.639072

**Authors:** Kiri Stern, Jin Vincent, Zofia E Taranu, Nathalie Fortin, Alexis Martel, Andrea Robbe, Romane Thaize, Mathieu Castelli, Stephen J. Beckett, Timothee Poisot, Angus Buckling, B. Jesse Shapiro, Nicolas Tromas

**Affiliations:** McGill University - Department of Microbiology and Immunology; Environment and Climate Change Canada; INRAE CARRTEL; Université de Savoie Mont Blanc; University of Maryland - Department of Biology; Université de Montréal - Départment des sciences biologiques; University of Exeter - Department of Ecology & Conservation; University of Maryland Institute for Health Computing; McGill Centre for Microbiome Research

**Keywords:** Cyanobacteria, phages, phage-bacteria interaction network, diversity

## Abstract

Phages – viruses that infect bacteria – are often seen as key players in bacterial community dynamics and the ecosystem services those communities support. However, much of our understanding of phage-bacteria interactions comes from in vitro studies, which provide limited insights into how these interactions occur in natural environments. In this study, we used cyanobacterial blooms as a model microbial community to evaluate the potential role of phages in driving cyanobacterial populations. These blooms are characterized by massive and usually rapid accumulation of cyanobacterial biomass and occur worldwide, threatening aquatic systems. The frequency and intensity of cyanobacterial blooms are increasing over time, likely due to human activities, and, until now, the role of predators like phages has remained unclear. We used a deep amplicon sequencing approach targeting the g20 capsid gene to profile the cyanophage community in eutrophic Lake Champlain over time, comparing their dynamics and diversity with bacterial communities and examining their associations. We evaluated whether phages simply followed bacterial dynamics, with limited impact on bacterial composition, or instead whether they played an active role in drving bacterial community structure. We found that phages exhibited similar dynamics to their potential bacterial hosts and shared environmental niches. However, we also observed strong differences specific to the phage community, such as inter-annual variation and an increase in Shannon diversity over time. Phage-bacteria and phage-cyanobacteria co-variance uncovered potential interactions that resulted in strongly modular networks. However, while the network structure remained modular, the composition of the modules differed significantly between environmental and temporal conditions. Lastly, viral phylogeny partially explained phage-bacteria co-variance, but only for a small proportion of bacterial ASVs. Our results indicate that phage-bacterial interactions are partly genetically structured and vary within modules across environmental conditions.

## INTRODUCTION

Microbial communities are composed of the most abundant and diverse organisms on Earth: Bacteria, Archaea, single-celled Eukarya, and viruses [1, 2]. Bacteriophages (or simply phages) are viruses that infect bacteria or arcahea [3]. Phages are the most abundant biological entities on Earth, outnumbering bacteria in most environments [3, 4] and often imposing significant bacterial mortality – from 20 to 40% in marine environments [5–7]. The interactions between phages and their bacterial hosts are therefore hypothesized to play an important role in the evolutionary dynamics and ecological structures of microbial communities. Phages are thought to drive bacterial population dynamics, altering intra- and inter-specific competition among bacterial hosts [6, 8–11], as described in the “kill the winner hypothesis” [12, 13] in which the fittest (most abundant) bacterial genotype is killed off by a specific phage, eventually to be replaced by a new winning genotype [14–17]. By this and other mechanisms, bacteria-phage coevolution tends to increase microbial diversity, for example by selecting for multiple modes of host resistance and increasing phage infectivity [18, 19]. Phages could therefore play an important role in driving bacterial dynamics and diversity [20, 21]. However, in some cases phages may merely track the growth and physiological states of their microbial hosts (17), with limited influence on community composition. A large part of this knowledge comes from simplified laboratory systems, and there is still a lack of understanding regarding the extent to which this will translate into infectivity in natural settings. Further *in-situ* studies are therefore required to understand the extent to which phages are “drivers” or “followers” of bacterial dynamics.

The structure of phage-bacteria interaction networks (PBINs) can help us understand the underlying ecology and evolution of phage-bacterial dynamics. PBINs have been studied experimentally by laboriously testing all pairwise combinations of phage and bacterial isolates [22]. Meta-analyses from soil [1] and oceans [23, 24] have revealed that PBINs tend to be non-randomly structured. The two most frequently observed structures are nestedness and modularity [25, 1]. Nested networks include both specialists (i.e unique host-phage pairs) and generalists (i.e phages that can infect multiple hosts). This structure is generally explained by a “gene-for-gene” infection model [26], where mutations provide bacteria resistance against recently evolved phages while maintaining previous phage resistance and host infectivity [1, 27]. By contrast, a modular network describes interactions that occur within distinct groups (or modules), rather than between groups [23]. Modular networks are explained by a “matching allele” infection model (also known as the Red Queen model [28] ) where each interacting organism evolves by losing their capacity to resist its ancestral partners [29]. Network analyses are therefore useful to identify co-evolutionary patterns and to quantify phage-bacteria interactions. However, one of the main weaknesses of most previously described PBINs is that phage and bacteria were collected at different sites from similar environments [1] and were maintained in high nutrient culture before infections were assayed in the lab. It is therefore unclear if and how the observed patterns can be generalized to phage-bacteria interactions in natural environments.

Co-occurrence networks provide a flexible analytical tool that could be useful to determine patterns from microbial communities in nature. Microbial interactions often drive co-occurrence relationships. For instance, taxa sharing the same niches dimension are more likely to co-occur, while competition may result in mutual exclusion. However, co-occurrence does not necessarily mean that taxa are directly interacting, and inferred networks could contain both true and spurious interactions [30]. To investigate microbial community dynamics, an appropriate scale of genetic resolution must be chosen. Ahlgren *et al.*, (2019) [31] described the importance of using very fine-scale, as opposed to coarse, higher taxonomic resolution for genotype analysis of network interactions between phages and different bacterial Amplicon Sequence Variants (ASVs). Fine-scale genotyping permitted the discovery of ecologically distinct populations of ASVs within marine cyanobacteria, which, in turn, lead to the identification of time-lagged predator-prey relationships with associated phages [32]. Studies have concluded that high genotypic resolution improves the analysis of microbial dynamics and that these fine scale observations clarify the role viruses have in shaping host microbial community structure [32–34].

Cyanobacteria have become a focus of attention due to their increasing occurrence in freshwater systems around the world [35, 36], causing harmful blooms and often producing cyanotoxins [37, 38]. Of the three described families of phages known to infect cyanobacteria (*Myoviridae*, *Podoviridae*, and *Siphoviridae*), a large majority belongs to the *Myoviridae* family [39, 40]. In this study, we used an amplicon sequencing approach on the viral capsid assembly gene (g20) [41], to target cyanophages from the *Myoviridae* family and evaluated how they might drive cyanobacterial communities.

Here, we present an 11-year time course study (2006-2016) of phage and bacterial communities in a large eutrophic North American lake, Lake Champlain, where cyanobacterial blooms are observed nearly every summer. In a previous study of this lake, we using 16S rRNA gene amplicon sequencing to identify *Microcystis* and *Dolichospermum* as the primary bloom-forming cyanobacteria [43]. Using bacteria community composition, we were able to predict the start date of bloom events with 78-92% accuracy. Blooms may end for various reasons, including abiotic environmental conditions and biotic factors such as phage predation or eukaryotic grazers of cyanobacteria. Here we combine amplicon sequencing of both bacterial and phage taxonomic marker genes to associate phage taxa with their hosts via co-occurrence networks, and to determine the extent to which phage and bacterial community dynamics are correlated over time. Our results highlight how phages are not simply drivers of bacterial community structure, arguing for a more nuanced view of phage-bacteria interactions in nature.

## MATERIAL / METHODS

### Sampling

Data sampling and pre-processing methods followed the same protocol outlined in Tromas, et al., (2017). 166 samples were collected from within the first meter of depth (photic zone) of Missisquoi Bay, Lake Champlain, Quebec, Canada (45°02’45’’N, 73°07’58’’W). Between 11 and 23 (median 15.5) samples were collected each year from 2006 to 2016, excluding 2014, between the months of March to October. 94 samples were taken from the littoral zone and 72 from the pelagic zone (Supplemental figures, Fig. S1). 0.2-μm hydrophilic polyethersulfone membranes (Millipore) were used to filter 50-250mL of lake water, depending on planktonic biomass density. Environmental data such as water temperature, average air temperature over one week, cumulative precipitation over one week, microcystin toxin concentration, and total and dissolved nutrients (phosphorus and nitrogen) were recorded during most sampling events as described in Tromas et al., 2017 [42].

### DNA extraction, purification, and sequencing

Phenol-chloroform purification and enzymatic lysis was performed to extract DNA from the 2006-2013 frozen filters, as per Fortin, et al., (2010) [43]. Samples were resuspended in 250μL of TE (Tris-Cl, 10mM; EDTA, 1mM, pH 8) and quantified with the PicoGreen dsDNA quantitation assay. DNA samples taken between 2015 and 2016 were extracted using PowerWater kit and quantified with NanoDrop. CPS1 and CPS2 primers were chosen as they target the viral capsid assembly gene (g20) and yields a fragment of 165bp [41]. This region was chosen due to its 80% specificity for marine water Cyanophages and amplification of *Myoviridae*, along with other phages. Previous studies have positively amplified cyanophages from freshwater system using this set of primers [44]. We therefore amplified the gp20 gene, then confirmed the library size by agarose gels and quantified DNA with a Qubit v.2.0 fluorometer (Life Technologies, Burlington, ON, Canada). Sequencing was performed at Genome Quebec on illumina MiSeq (250bp paired-end reads). We acknowledge that while the primers were designed to target cyanobacterial phages, the infection capacity of these phages is still unknown. In other words, these phages might also infect other bacteria.

### Sequencing data analysis

In order to remove primers, Cutadapt [45] was used. Sequences were then preprocessed using Mothur (version 1.35.1) [46] to trim sequence lengths between 120-200 base pairs and to remove outliers. Using the DADA2 (library version 1.14.1; [47]), workflow, single-nucleotide resolved ASVs were inferred. Forward and reverse read pairs were trimmed and filtered, dereplicated, chimera-checked, and merged using standard parameters (http://www.metagenomics.wiki/tools/16s/dada2). 4502 ASVs from the 9,698,187 sequences processed through DADA2 were obtained and stored as a Phyloseq (R package version 1.44.0;[48]) object, ranging from 23 to 130,512 reads per sample, with a median of 59,848. Six samples with fewer than 3000 sequences were removed from the ASV table using Phyloseq. Bacterial ASVs were previously inferred and reported in Tromas, et al., 2017 [49]; using the same samples.

### Spatio-temporal analysis

To describe the viral community’s composition of conditionally rare taxa (CRT), low abundant taxa, and seasonal taxa we followed Shade, et al., (2014) [50] R script for detecting CRT in a temporal microbial community dataset using the default values. To calculate the number of low abundant taxa, we filtered out all ASVs whose sum across all samples was equal to zero and identified the number of ASVs whose relative abundance across samples was below the set the detection threshold of <0.0005 as defined by Shade, et. al., (2014) [50]. Finally, to determine the total number of seasonal viral ASVs, we filtered out all ASVs whose sum across all samples was equal to zero and counted the number of produced Lomb-Scargle periodograms generated by the R package Lomb (version 1.2; Ruf, T., 1999 [51]). Lomb-Scargle periodograms are a commonly used method to evaluate seasonal events due to its proficiency in detecting weak trends in unevenly sampled time series data [51].

To investigate the importance of spatio-temporal variables on viral diversity, we calculated the beta diversity using the square root of the Jensen-Shannon divergence (JSD) distance between the different sites (Pelagic and Littoral) and temporal variables (Day of the Year, Weeks, Months, Seasons and Years). The permutational multivariate analysis of variance (PERMANOVA) test using the adonis2() function in the library vegan (version 2.6-4; [52]) was used to determine whether these variables were significant enough to explain the variance. As this approach can be sensitive to dispersion, we also tested for dispersion by performing an analysis of multivariate homogeneity (PERMDISP, [53]) with the permuted betadisper() function.

### Comparison of alpha diversity between phage and Bacteria

The composition of the viral ASVs was first described by selecting the 20 most abundant viral ASVs from both locations (*i.e.* the littoral and pelagic zones). We used microeco R package to generate these compositional plots [54]. We then characterized the diversity of viral ASVs by calculating total richness and Shannon diversity. Littoral and pelagic sites were analyzed separately (Supplemental Materials Figure S.1).

To characterize alpha diversity, we calculated total richness using Breakaway (package version 4.6.16 ; [55]) due to its robustness in estimating microbial biodiversity and incorporation of standard errors. Breakaway was designed specifically to consider the structure often observed in microbial datasets characterized by a high proportion of rare species, “singleton” counts, and very few abundant species [55]. The Shannon diversity was measured using the estimate_richness() function in the library Phyloseq in R. Viral ASVs richness and Shannon diversity was plotted against each sample, years, bloom, as well as across sampling depth. While richness could be driven by sampling depth and therefore making any observations difficult to interpret, Shannon diversity was not associated with depth variation.

Finally, a cross-correlation function (CCF) analysis was conducted to assess the temporal relationship between phage, bacterial and cyanobacterial Shannon diversity using the ccf() function of the tseries R package [56]. No specified maximum lag was specified (lag.max=NULL) to allow for an automatic selection of the lag range based on the data length. Both correlation and covariance types were examined, though only correlation results are reported here.

### Associations with environmental variables

To examine the relationships between taxa and environmental factors, we performed a redundancy analysis [57]. This method identifies the linear combination of explanatory variables (the matrix of abiotic environmental data) that best accounts for the variation in the response matrix (the viral ASVs table). We first Hellinger-transformed the viral ASVs table [58], and standardized the explanatory (environmental) table using z-scores with the function decostand(x, method = ’standardise’) to account for differing units among the environmental parameters. RDA was performed using the rda(scaling = 2) function from the R vegan package and we used the anova.cca() function from the R vegan package to evaluate the significance of constraints.

### Viral-Bacterial dynamics

We compared the bacterial and phage dynamics dynamics over time (day of year; DOY) using a Latent Dirichlet Allocation (LDA) machine learning [59] with methods adapted from Christensen et al. (2018) [60]. We first calculated the relative abundances of each ASV by normalizing its read count against the total read count for each sampling date. To reduce the dimensionality of the data matrices and capture changes in community composition over the seasons, we employed three approaches. First, we constructed a virus-host interaction network based on Spearman correlation coefficients, retaining interactions with edge probabilities exceeding 0.65. This reduced the virus matrix from 550 ASVs to 244 and the host matrix from 1059 to 95 ASVs. Secondly, we applied the dropspc() function from the R package labdsv [61], setting minocc=1 and minabu=0.01 to remove taxa with less than 1% relative abundance or less than one occurrence through time. This further reduced the virus matrix to 123 ASVs and the host matrix to 76 ASVs. Following these initial filtering steps, we aggregated ASV-level information into community-types, further reducing dimensionality, using LDA approach. To mitigate the impact of highly variable read counts on LDA estimation, each ASV’s relative abundance was scaled by 100 and rounded down to the nearest integer [62, 63]. The number of community-types was determined by evaluating model AIC across varying numbers of types and selecting the model with the lowest AIC.

Once dimensionality reduction and identification of community-types were completed, we analyzed changes in community composition using a Bayesian change point model tailored for historical time series [60, 64]. This approach divides the time series into segments at change points, with each segment characterized by its own set of parameters. The Bayesian framework integrates prior knowledge and updates posterior distributions as new data becomes available, employing Markov Chain Monte Carlo (MCMC) sampling to derive the posterior distribution of model parameters and change point locations. Unlike methods assuming a static process, the Bayesian change point model is adaptable to shifts or tipping points in the data over time. Outputs include histograms depicting sampled change point locations, highlighting where changes are most likely.

### Correlations and covariance

To understand as well the intra-relationship between viral ASVs and the inter-relationship between viral and bacterial ASVs, we calculated the correlation and covariance coefficients between ASVs. The viral and bacterial phyloseq objects were filtered to remove taxa not seen more than once in at least 10% of the samples to protect against ASVs with small means and trivially large coefficients of variation.

The strength of phage-bacteria covariance was inferred using the SParse InversE Covariance estimation for Ecological Association Inference (SpiecEasi; [65] ; version 1.0.7) R package. SpiecEasi determines cross-domain interactions from merged ASV count tables and can predict direct associations using conditional independence, as well as compute negative covariance[66]. Covariance analysis was therefore performed within viral (A) ASVs only, (B) between viral ASVs and bacteria, (C) between viral ASVs and Cyanobacteria, and (D) between viral ASVs and *Microcystis*/*Dolichospermum* ASVs.

Neighborhood selection (mb) was used to infer the graphical model and parameters for each viral covariance analysis were updated to minimize the difference between the output and stability threshold (0.05). We filtered the networks (edge-based confidence filter > 0.7) to keep the most stable edges using getOptMerge function. The igraph package ([67]; version 1.2.6) was used to visualize ASV associations and find community structure by grouping clusters into modules using the fast greedy modularity optimization algorithm in the cluster_fast_greedy() function.

### Relationship between phage phylogeny and phage-Bacteria co-occurrence

We asked if closely related viral ASVs have a higher chance to co-occur and if closely related viral ASVs have higher probability to co-occur with similar Bacteria (*i.e.* potentially sharing the same hosts). We first estimated the genetic distance between each pair of ASVs using DistanceMatrix R function (Decipher R package, version 2.0.2; [68]. To determine if closely related viral ASV co-occur with similar bacteria, and thus potential host, we calculated the absolute difference of SpiecEasi covariance between each viral ASV (Xi) and a specific taxon T. We used a similar strategy as described in Tromas et al., 2020 [69], where we calculated |Δr| = | Co(X1, T) -Co(X2, T) | with Co defined as the SpiecEasi covariance between Xi and T. For bacterial ASV, we estimated the relationship between |Δr| and the distance between the given viral ASV (X1 and X2). A positive correlation would indicate that more closely related phages would have more similar correlations with potentially interacting bacteria. Finally, we conducted a permutation test to assess potential bias in the method arising from the data structure. For each significant bacterial ASV, we estimated the rate of false-positive correlations by resampling viral ASV genetic distances with replacement while randomizing the association between genetic distance and |Δr|. We performed 1000 permutations test and recorded the proportion (p) of permutations yielding a larger correlation than observed, after adding a pseudocount of 1 to both the numerator (the number of permutations yielding a larger correlation than observed) and the denominator (the total number of permutations).

### Nestedness and modularity

The degree of nestedness of the bipartite phage-bacteria interactions was measured in R using FALCON software ([70]; version 1.0). FALCON offers the option to choose between six different nestedness measures (Spectral radius (SR), the measure of Johnson, Domínguez-García, & Muñoz (JDM), discrepancy (BR), Manhattan distance (MD), nestedness based on overlap and decreasing fill (NODF), and nestedness temperature calculator (NTC)) and five null models (Swappable-Swappable (SS), Fixed-Fixed (FF), Cored-Cored (CC), Degreeprobable-Degreeprobable (DD), and Equiprobable-Equiprobable (EE)) for nestedness in binary networks; and two nestedness measures (weighted-NODF and SR) and four null models (binary shuffle, conserve row total (CRT), conserve column total (CCT), and row columns totals average (RCTA)) for weighted networks. Descriptions for each measure and model can be found in the FALCON manuscript.

We ranked rows and columns by degree to maximise nestedness, selected the NODF measure to calculate the nestedness score, and used the DD null model as the comparison. In studies of interaction networks, SS, FF and DD are commonly used. SS and FF keep the same number of phage and bacteria; SS places the same number of interactions randomly; FF places them such that the row and column sums of the matrix are the same as in the initial network. DD uses the initial matrix as a framework and assigns elements probabilistically according to the network degree distributions. As the SS model is very lenient (i.e. it is very easy to find networks less nested than the original) and the FF model is potentially quite strict (can be hard to find networks less nested), we opted for the DD null model.

Beckett (2016) [71] released algorithms to maximize weighted modularity in bipartite networks: LPAwb+ and DIRTLPAwb+. Due to its efficiency with large datasets, we used the LPAwb+ to calculate the modularity score. We then extracted modules information to compare modules composition.

## RESULTS

### Viral and bacterial community structures both vary seasonally but diverge over yearly time scales

In Tromas et al., (2017) [42], we conducted an analysis of the bacterial community from an eight-year time course (2006 – 2013) of Lake Champlain. We found that community structure changes cyclically over the course of a year, with ‘Week of the year’ and ‘Day of the year’ explaining most of the variation. Here, we describe an overlapping but longer time series (2006-2016) of the viral community in Lake Champlain, focusing on cyanophages from the *Myoviridae* family. We first compared the time scales explaining most of the variation in bacterial and viral communities. Most of the variation in both viral and bacterial communities was explained by the week of the year, with 43% and 44% of the variation respectively explained, suggesting clear seasonal patterns (**Table 1**). However, nearly twice as much variation (27%) in the viral community was explained by the year of sampling, compared to bacterial communities (13%). This difference was not due to differences in dispersion between bacterial and viral communities (betadisper *p* > 0.05). As described previously, bacterial communities were significantly impacted by blooms (*R^2^* = 9%) but the effect of blooms on viral communities, while significant, was much smaller blooms (*R^2^* = 2%; **Table 1**). Together, these results suggest that bacterial and viral communities both vary seasonally but differ in their response to blooms may diverge over longer, yearly time scales.

**Table 1.** Spatio-temporal analysis of the viral and bacterial community structure. PERMANOVA tests were applied to Jensen-Shannon divergence (JSD) of the relative abundances of viral or bacterial amplicon sequence variants (ASVs).

| VARIABLES | PERMANOVA | PERMANOVA | PERMANOVA | PERMANOVA |
| --- | --- | --- | --- | --- |
|  | R<sup>2</sup> | P-VALUE | R<sup>2</sup> | P-VALUE |
|  | Viral ASV |  | Bacterial ASV |  |
| **SITE** | 0.28% | 0.302 | 0.64% | 0.904 |
| **BLOOM** | 2.35% | 0.004 | 9.41% | 0.001 |
| **DOY** | 37.28% | 0.001 | 38.46% | 0.001 |
| **WEEK** | 43.55% | 0.001 | 43.80% | 0.001 |
| **MONTH** | 21.49% | 0.001 | 21.31% | 0.001 |
| **SEASON** | 9.37% | 0.001 | 11.15% | 0.001 |
| **YEAR** | 26.89% | 0.001 | 13.42% | 0.001 |

To explore this intra and inter annual variability, we visualized the dynamics of the most abundant viral ASVs and confirmed their temporal variation. For example, viral ASV4 was present only some years (e.g. 2006) while viral ASV1 was present every year, but not at the same time of year (Supplementary Figure S1). We then compared the proportion of the seasonal ASVs (*i.e.* those with a recurrent peak of abundance during same season) and rare ASVs across the viral and bacterial datasets. Seasonal ASVs were more abundant the in viral community (2%) compared to the bacterial community (1%). The proportion of conditionally rare ASV, defined as ASV having a bi-modal abundance over time with one or few peaks of abundance [50] is also more abundant in the viral community. The most important difference lies in the rare biosphere that is more abundant in bacterial (68%) than viral community (39%) (**Table 2**).

**Table 2.** Viral and bacterial communities contain similar proportions of rare and conditionally rare taxa (CRT).

|  | Viral ASV | Bacterial ASV |
| --- | --- | --- |
| **Total CRT (bimodal)** | 87 \| 2.7% | 110 \| 1.7% |
| **Total low abundance (<0.005)** | 1241 \| 38.9% | 4224 \| 68.4% |
| **Seasonal ASV** | 70 \| 2.2% | 64 \| 1.0% |
| **Total ASV** | 3185 | 6168 |

Other variables were tested to explain variation in viral community structure. As observed previously for the lake bacterial community, ‘Site’ was not significant (p-value = 0.302) indicating that the viral community between Littoral and Pelagic sites were not different. Water temperature and dissolved nitrogen explained variation in both phage and bacterial community structure, while several others factors such as total phosphorus, total nitrogen and precipitation were strong drivers of bacteria but not viral communities (Supplementary Figure S2). The redundancy analysis explained relatively little variation in the viral community (*R^2^* = 14%, *p* = 0.001), compared to a moderate amount in the bacterial community (*R^2^* = 19%, *p* = 0.001). These results further indicate the distinct dynamics and drivers of viral compared to bacterial communities.

### Phages and Bacteria shared similar dynamics shifts across years

To evaluate the potential interactions between phages and their host, we compared their dynamics by characterizing the temporal succession of viral and bacterial communities. We applied a Latent Dirichlet Allocation (LDA) approach (Christensen et al. 2018) to identify “topics” or types within each of the two communities that tended to co-occur in set proportions. By viewing how these different communities varied over the time (DOY), LDA model selection identified 14 bacterial and 15 viral community-types as optimal. We then analysed dynamics changepoint to highlight when, if any, major changes occurred among the identified topics. The change point analysis identified similar significant shifts at Day of Year (DOY) 189 (first week of July, corresponding on average to the beginning of bloom) and approximately DOY 260 (mid-September, corresponding on average to the end of bloom) for viral and bacterial community, indicating temporal potential correlations. Interestingly, the most important shift for phages occurred during a bloom event (DOY 220), after the most important shifts in bacterial community-types. Most community-types were dominated by just one of very few ASVs (e.g., the bacterial community-type 3 was composed of 93% ASV 55 which was classified as *Synechococcus*), (Supplementary Figure S3 – S4). Other community-types were composed of bloom-forming Cyanobacteria (*e.g Microcystis* and *Dolichospermum*) and photosynthetic eukaryotic organisms (*i.e.* classified under “Chloroplast”). This result suggested that across years, phage and bacteria shared shifts in their dynamics.

### Viral Shannon diversity increases over years and is correlated with cyanbacterial diversity at varying time lags

We next evaluated how viral alpha diversity varied over time, and if it was related to bacterial diversity. We calculated the viral ASV richness and Shannon diversity within each sample and found that both increased significantly over years (Figure 1, Supplementary Figure S5). While richness was confounded by sequencing depth (linear regression, R^2^=20%, *p*<0.001), Shannon diversity was not (Supplementary Figure S6, linear regression, R^2^∼0*, p* >0.05). The increasing Shannon diversity over our decade-long sampling could be a due to a combination of increased richness and increased evenness in the viral community. In contrast, bacterial richness and Shannon diversity decreased significantly over years (Figure 1). However, this result might be potentially biased as richness and Shannon diversity increased over sequencing depth (linear regression, R2=60.7%, p<0.001; R2=7%, p<0.01). Consistent with relative lack of viral community changes during blooms, we did not find any difference in viral community alpha diversity between bloom and non-bloom conditions (Supplementary Figure S7, Wilcoxon test p-value>0.05). Bacterial and viral Shannon diversity values were not significantly correlated (*p* > 0.05 for both total bacteria and for cyanobacteria only). This lack of correlation further highlights the distinct dynamics of bacterial and viral communities, although we cannot exclude technical explanations (*e.g.* different levels of genetic resolution in viral and bacterial marker genes) for the lack of correlation.

**Figure 1.**
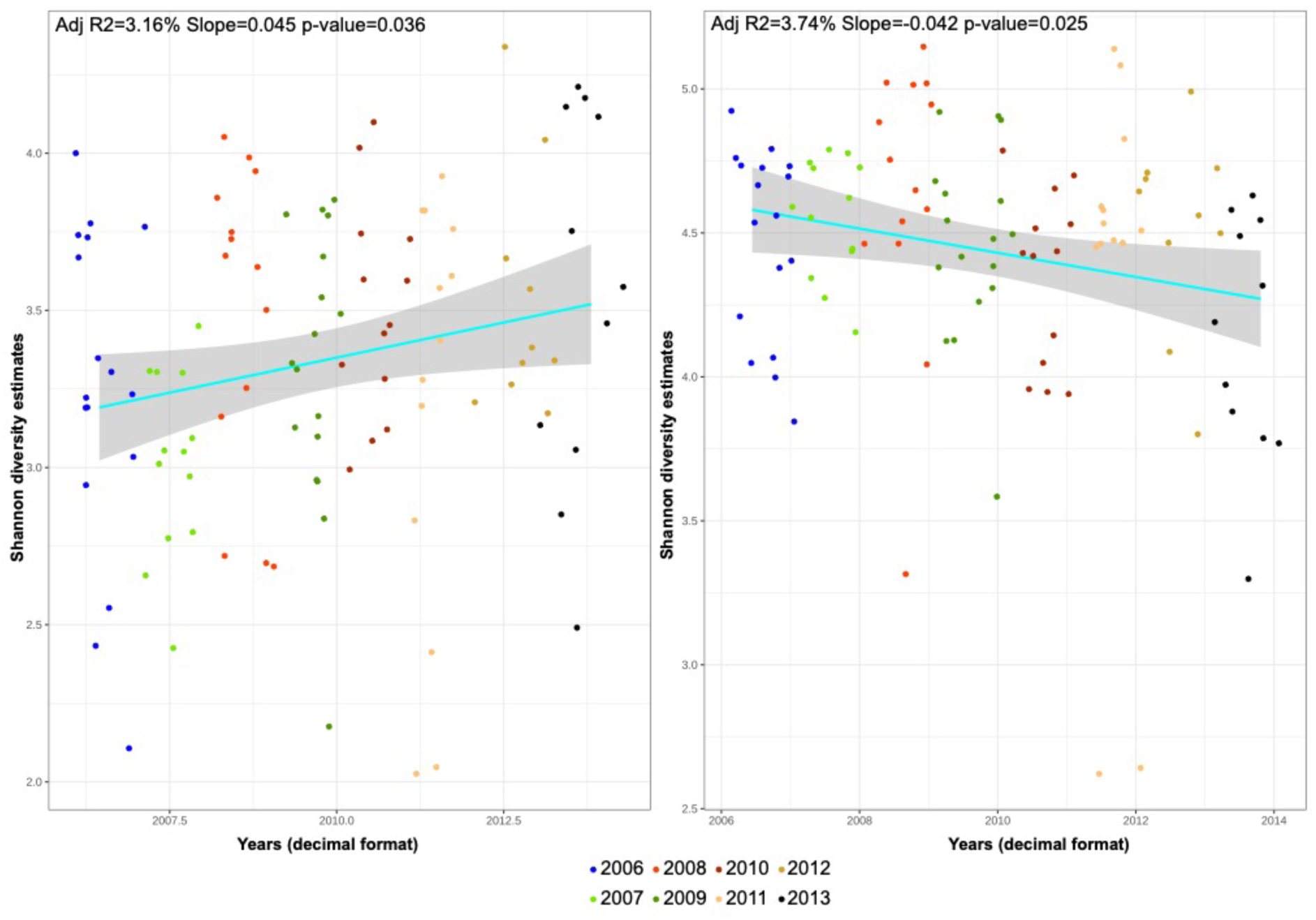
Shannon diversity across years.

Despite the lack of correlation between viral and bacterial diversity from the same sample, it is possible that viral diversity could impact future levels of bacterial diversity, or vice versa. For example, in “kill the winner” a dominant bacterial genotype is driven to lower abundance by a specific phage, increasing bacterial Shannon diversity, and potentially generating a pattern of low phage diversity being predictive of increased bacterial diversity on a certain time scale. To identify such patterns, we calculated time-lagged correlations between bacterial and viral diversity. There were no significant correlations between bacterial and viral Shannon diversity at any time lag. However, significant correlation between cyanobacterial and viral Shannon diversity were observed at several lags (Figure S9). Significant negative correlations (-0.2; -0.24; -0.26) at lags of -13, +1 and +2 and a positive correlation (0.24) at lag +7 time points were observed. This indicates that changes in cyanobacterial diversity precede changes in phage diversity by 13 time points. Phage diversity then leads to a decrease of cyanobacterial diversity with a delay of 1-2 time points. Finally, phage diversity leads to higher cyanobacterial diversity after 7 time points (Supplementary Figure S8). While it is difficult to precisely interpret the biological meaning of these patterns, they imply that cyanobacteria may be drivers of phage diversity on some time scales, while the reverse may be true on other time scales.

### Phage-bacteria interaction networks

To our knowledge, there are very few studies that have described phage-bacterial interaction networks (PBINs) across time and environmental conditions, in natural settings. We constructed PBINs for different combinations of communities: phage-Cyanobacteria, or phage-*Dolichospermum*-*Microcystis* (Supplementary Figure S9-10) – the two most dominant bloom-forming cyanobacterial genera in Lake Champlain [72]. We also constructed separate networks from samples with different nutrient conditions, from different seasons, and during blooms. All the resulting networks were highly modular and not nested (Supplementary Tables 2) suggesting specific phage-bacterial interactions within modules [1]. We found that a very large majority of the modules are composed of only one cyanobacterial ASV and from one to seven viral ASVs (∼2.5 on average). Normalized and realized modularity values were higher for ‘Eutrophic’ and ‘bloom’ networks but not necessarily for ‘summer’. We next evaluated how the module composition varied across conditions (Supplementary Figures S11&12, Supplementary Tables 3&4). The number of ASV composing the modules was reduced during bloom and eutrophic conditions (1 cyanobacterial ASV for ∼1.55 viral ASVs) compared to no bloom and mesotrophic condition (1 cyanobacterial ASV for ∼2.55 viral ASVs). We also found that nodes and edges were highly dissimilar between mesotrophic and eutrophic conditions, as well as between bloom and non-bloom conditions (Jaccard dissimilarity > 0.7). Thus, while the network structure always remains modular, the composition of phage, bacteria, and their interactions within modules changes. We repeat the analysis on the two most dominant bloom-forming Cyanobacteria - *Microcystis* and *Dolichospermum* – and found similar results with different co-occurrent networks across conditions (Figure 2), a structure that remained modular (bloom network realized modularity = 0.76, non-bloom network realized modularity = 0.96) with modules that are not similar across bloom conditions (Supplementary Tables 5). For example, *Microcystis* ASV 7 interacts with three viral ASVs in the global PBIN, but it interacts with three different ASVs in the bloom-specific network (Figure 2, Supplementary Figure S10). This highlights how distinct modules of interactions can arise during different environmental conditions, or over time.

**Figure 2.**
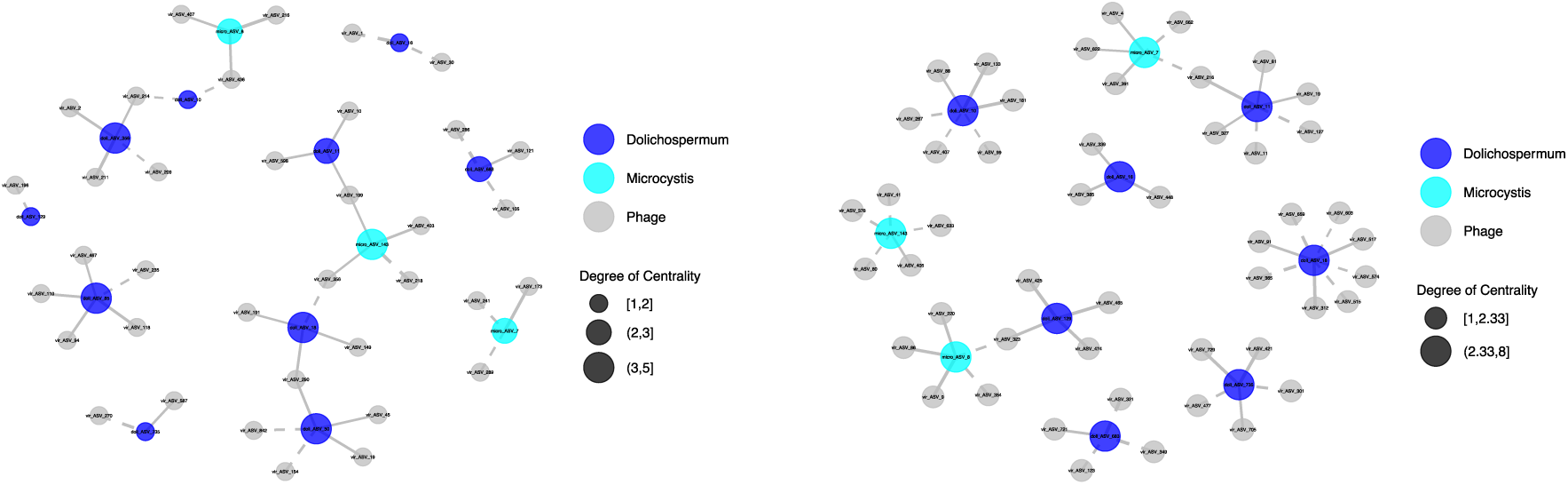
Phage-*Microcystis*/*Dolichospermum* co-variance network during bloom (left) and no bloom (right). Solid and dashed lines correspond to respectively positive and negative co-variance.

### Closely related phages tend to covary with similar bacteria

To further interrogate the structure of the inferred PBINs, we asked if more closely-related phages tend to co-occur with each other, and with similar bacteria. We found that more closely-related phages (in the ranges of 50%-100% and 90%-100% nucleotide identity in g20) had a tendency to co-vary with one another (Supplementary Figure S16). However, such increase in co-variance was not observed in reduced range of 95%-100% (Adjusted R-squared: -0.006 F-statistic: 0.2402 on 1 and 125 DF, p-value: 0.6249). We then performed a pairwise analysis to examine the relationship between the genetic distance between each pair of viral ASVs and |Δr|, a measure of the similarity of viral ASV co-variance with bacterial ASVs (Methods). To reduce false positive and noise, we only considered bacterial ASVs that co-varied significantly with at least five different viral ASVs. Of 798 bacterial ASVs meeting this criteria, the correlation between |Δr| and genetic distance was significant for 98 of them. A permutation test showed that for these 98 bacterial ASVs, there is significantly more ’signal’ in the data than expected by chance (Supplementary Figure S17). 71.5% of these significant correlations were positive relationships between co-occurrence and genetic distance, *i.e.* a higher proportion of more closely related viral ASV tend to co-occur with similar bacterial taxa. Together with the PBIN analysis, this result indicates that phage-bacterial interactions are partly genetically structured and varied within modules across environmental conditions.

## DISCUSSION

### Potential caveats

In this study, we used a set of primers that were designed for marine cyanophage and were previously tested on freshwater Cyanobacteria with different success [44, 73]. We acknowledge that these primers might not target all the cyanophages present in this freshwater system. We decided to include cyanobacterial ASVs taxonomically classified as ‘Chloroplast’ representing potentially photosynthetic eukaryotic organisms. If cyanophage should not target these micro-organisms, their dynamics were still interesting, especially as they might compete with Cyanobacteria. Finally, we used a co-variance network as a proxy for an interaction network, acknowledging that co-variance does not necessarily imply interactions.

### To some extent, phages might follow the same dynamic as Bacteria

Our results brought some evidence that phages could follow their potential host dynamics [74]. We found that viral community was explained by several similar temporal variables (e.g. ‘weeks’ and ‘Months’) that also explained bacterial community (Tromas et al., 2017); and also similar environmental variables, such as dissolved nitrogen and water temperature. LDA analysis showed that phages and cyanobacterial dynamics had similar changing points in their dynamics, suggesting potential correlation of dynamics across the time course. This results mostly concerned specific cyanobacterial subcommunities, including the most dominant bloom forming Cyanobacteria, photosynthetic eukaryotic organisms and *Synechococcus* ASVs. Finally, we found that viral diversity was also correlated to Cyanobacterial diversity with multiple different time-lags. Overall, these results showed that the dynamics and diversities of phage and Cyanobacteria are related and might support the hypothesis that phages may simply be driven by fluctuations in microbial densities, which are known to be influenced by seasonal changes and environmental conditions [74, 75].

### Phage might drive Bacteria

However, we also found difference between phages and bacterial dynamics. We observed differences with more rare and conditionally rare taxa within phage community. Shade, et al., (2014), proposed that rare taxa help maintain community stability by “acting as a reservoir that can rapidly respond to environmental changes”. These rare phages could, therefore, serve as such a reservoir. We found that temporal variables such as ‘Years’ was a strong explanatory variables, suggesting that viral community could vary intra-annually but also inter-annually which is different to what we previously observed with the bacterial community. We also found that Shannon diversity increased over years suggesting that phages might evolve to evade resistant bacteria, while unevolved phages might be retained. Finally, we found that ‘bloom’ poorly explained viral community variation and did not impact viral diversity. This result suggests that high disturbance in microbial community did not drive strong changes in viral community. This result is not what we expected as host density is generally a factor associated with phage dynamics [76]. A potential explanation of such observation is that the majority of phages selected by our g20 primers did not target the two most dominant bloom forming Cyanobacteria present in this lake (e.g. *Microcystis* and *Dolichospermum*). While previous studies [73, 77] showed that these primers, initially designed for marine cyanophage, successfully targeted freshwater cyanophages [44]; other freshwater viruses might be amplified. On the other side, LDA analysis showed that a significant changing points within viral community happened generally during bloom. As diversity is not impacted, we could hypothesized that viral subcommunities increased and other decreased in abundance during bloom, explaining such shift in community dynamic.

Overall, our results show that phage dynamics and diversity are in some ways different and decoupled from the putative cyanobacterial hosts. While there is some evidence that hosts could be drivers, and some evidence that viruses could be drivers; our observations showed that phage-bacterial interactions can be complex and could vary across time or environmental conditions.

### Phage-Bacteria interaction network

To explore these variations, we used co-occurrence network as a proxy for phage-bacteria interactions and found that across this 11-years time course, the network was highly modular. Modular networks are generally associated with fluctuating-selection dynamics and were previously found in low-nutrient environments [78]. From modular network, we extracted modules information and found that modules composition was mostly a one-to-one (*i.e.* on average, one viral ASV co-occurred with one specific cyanobacterial ASV) suggesting highly specific interactions. To evaluate if the composition of these modules changed across nutrient conditions and bloom conditions, we re-built independent phage-Cyanobacteria networks and re-calculated modularity. We still found highly modular networks across these conditions (Supplementary Table 3) but with different modules and edges composition. This result suggests that nutrient resources and host density is not directly driving – at least in this lake – the ecological and evolutionary responses of the cyanobacterial population to phages. However, the fact that modules composition varied across conditions showed that the potential interactions – within modules – between phage and bacteria could be driven by other variables, such host genotype. Previous studies [79] have shown that host phylogeny could be used as predictor of the susceptibility of bacterial strains to phage infection. Here, we used viral phylogeny to estimate the impact of potential host phylogeny and found that – for certain ASVs at least – closely related viral ASV have a higher chance to co-occur with the same Bacteria.

## Conclusion

In this study, we compared viral and bacterial dynamic over a decade using an amplicon sequencing approach. We targeted cyanophages from the *Myoviridae* family to evaluate if they could drive the cyanobacterial bloom that Lake Champlain suffered from mostly every year. While viral and bacterial communities were influenced by similar temporal variables, environmental parameters, and shared dynamic change points, they also exhibited significant differences. The structure of the viral community was composed of more rare ASVs and their diversity increased over years. Moreover, the viral community was weakly explained by bloom and the viral diversity was not impacted, despite the increase of cyanobacterial density. Finally, co-variance networks were used as proxi for phage-bacteria interaction networks. While co-variance between two ASVs does not imply interaction, it suggests a higher likelihood of one. We generated different co-variance networks between phage and Bacteria, Cyanobacteria, and the two most dominant bloom forming Cyanobacteria *Dolichospermum*-*Microcystis*. All networks were significantly modular and not nested. When we generated networks within specific conditions (*e.g.* bloom/no bloom or mesotrophic/eutrophic) networks kept their modular structure with strong changes in modules composition.

## Data availability

Raw sequence data have been deposited NCBI GenBank under BioProject number PRJNA1212704

## Acknowledgment

We thank everyone who participated in sampling, data collection and analysis, with special thanks to Alberto Mazza and Miria Elias. This work was supported by grants from NSERC Discovery Grant to BJS.

