## Supplementary_figures_tables for "Deciphering the dynamics of Cyanobacteria-Phage in a natural lake: Insights from a decade-long investigation"

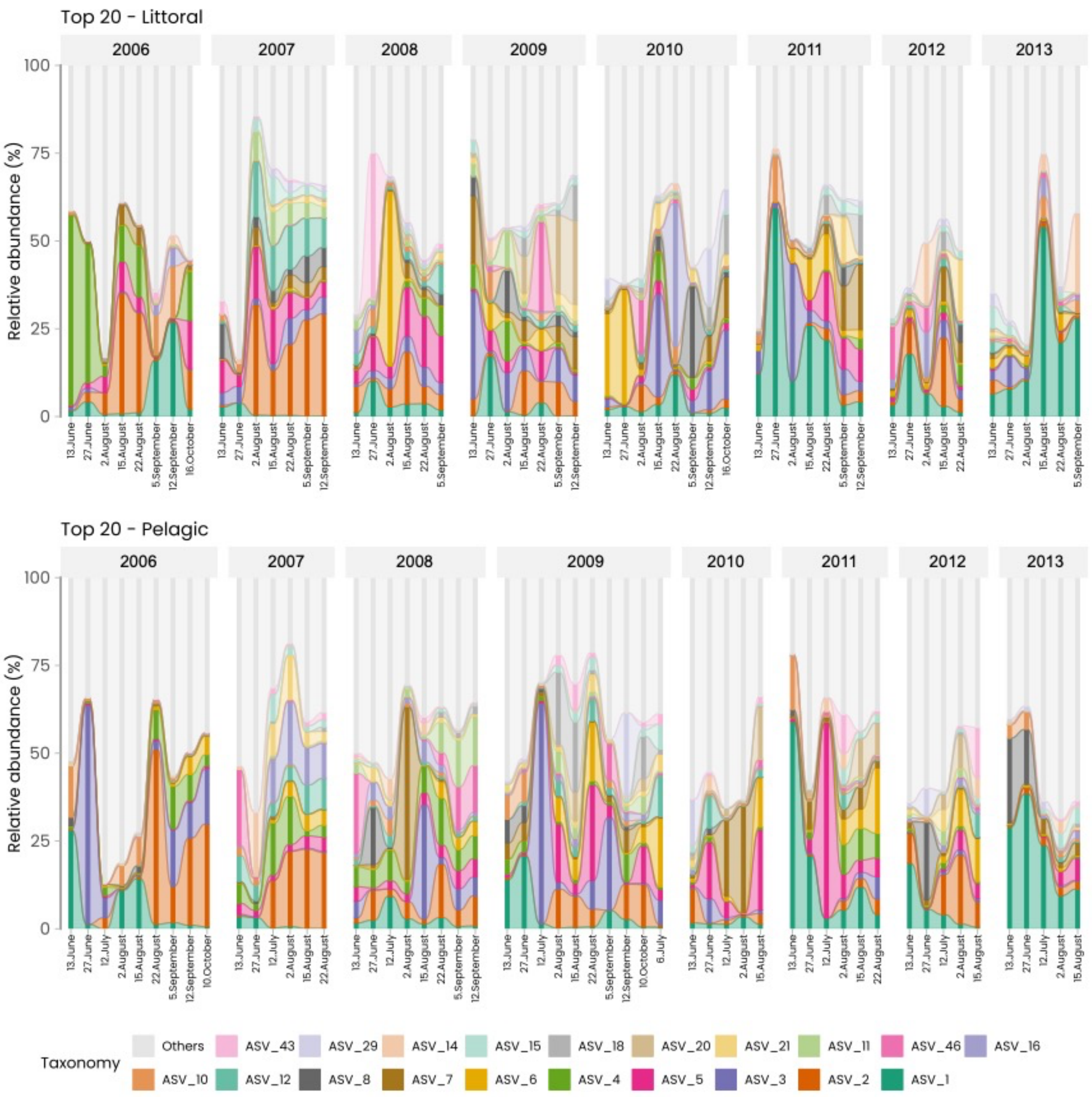


***S1. Relative abundance of the (top) Littoral ASVs and (bottom) Pelagic ASVs by date.***
*The relative abundance of the top 20 viral ASVs from the pelagic and littoral zones are shown.*

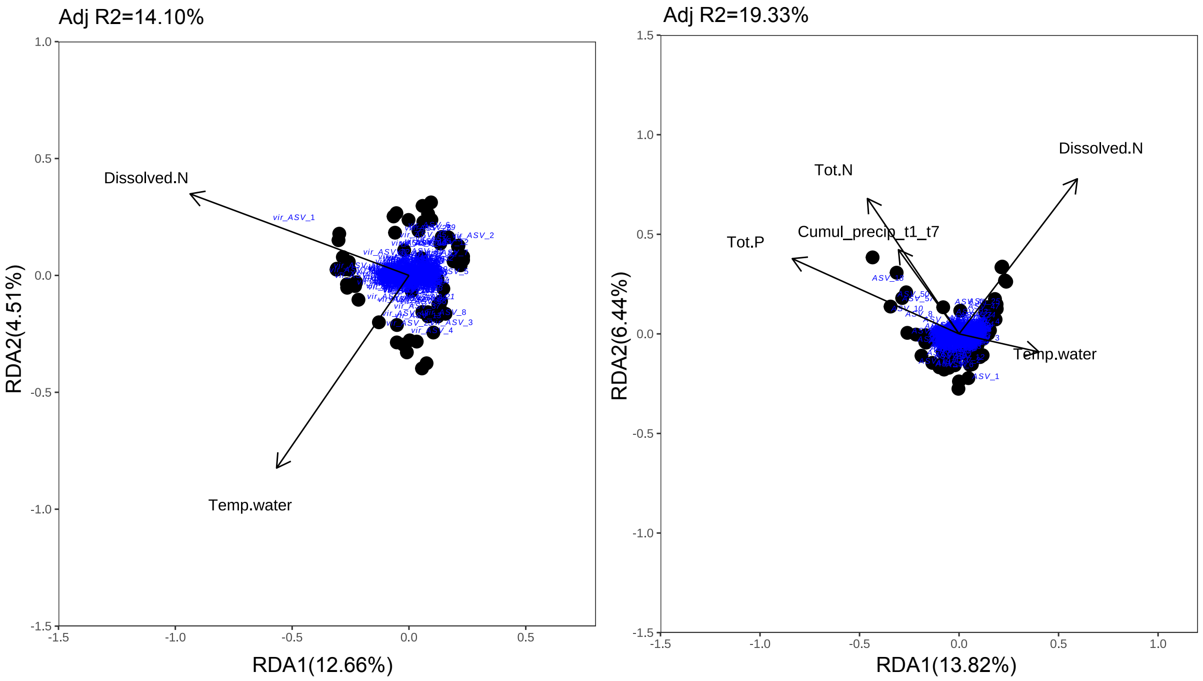


**S2. Phage (left) and bacterial (right) community dynamics are explained by different environmental variables.** A redundancy analysis was performed on selected environmental variables from ordiR2step function. NA were removed from the environmental variables table and the final number of observations for phages and bacterial community was 57 (initial table was composed of 108 samples). We then used an ANOVA permutation test for Constrained Correspondence Analysis (CCA) and found that both models were significant (p-value = 0.001). Both axes were also significant. Adjusted $R^2$ = 14.10% for phages and 19.33% for Bacteria.

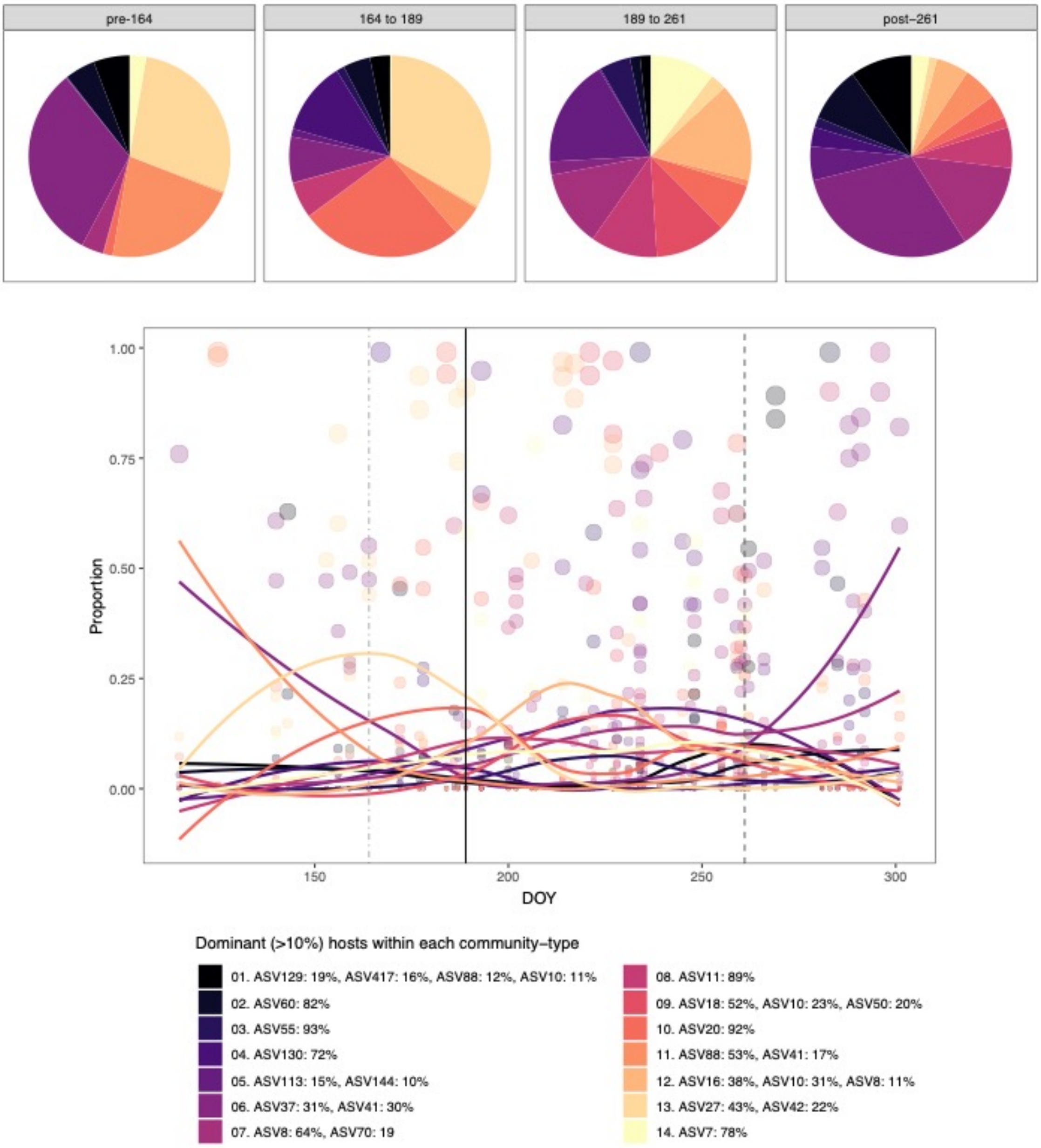


### S3. LDA bacterial communities dynamics

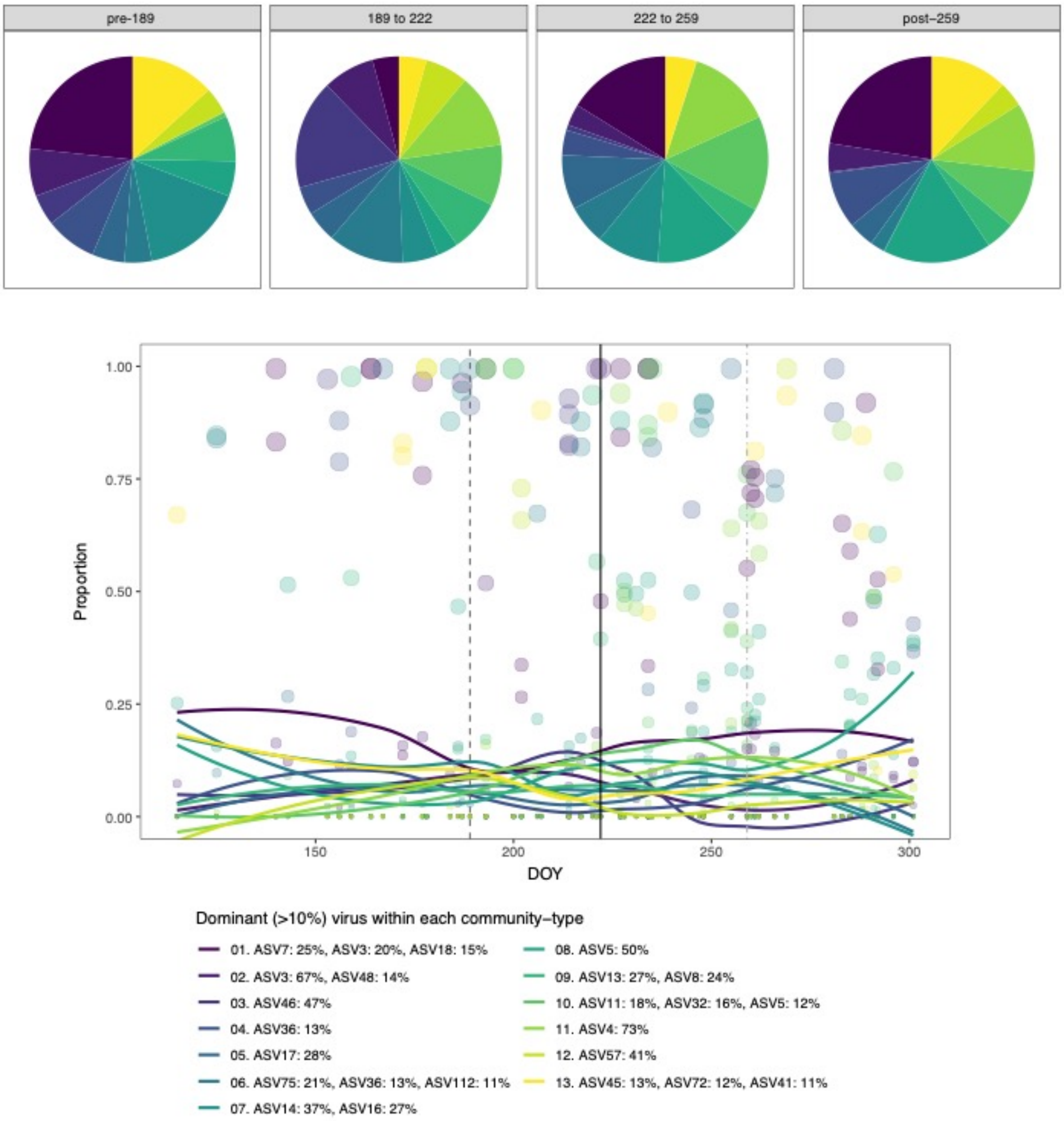


### S4. LDA viral communities dynamics

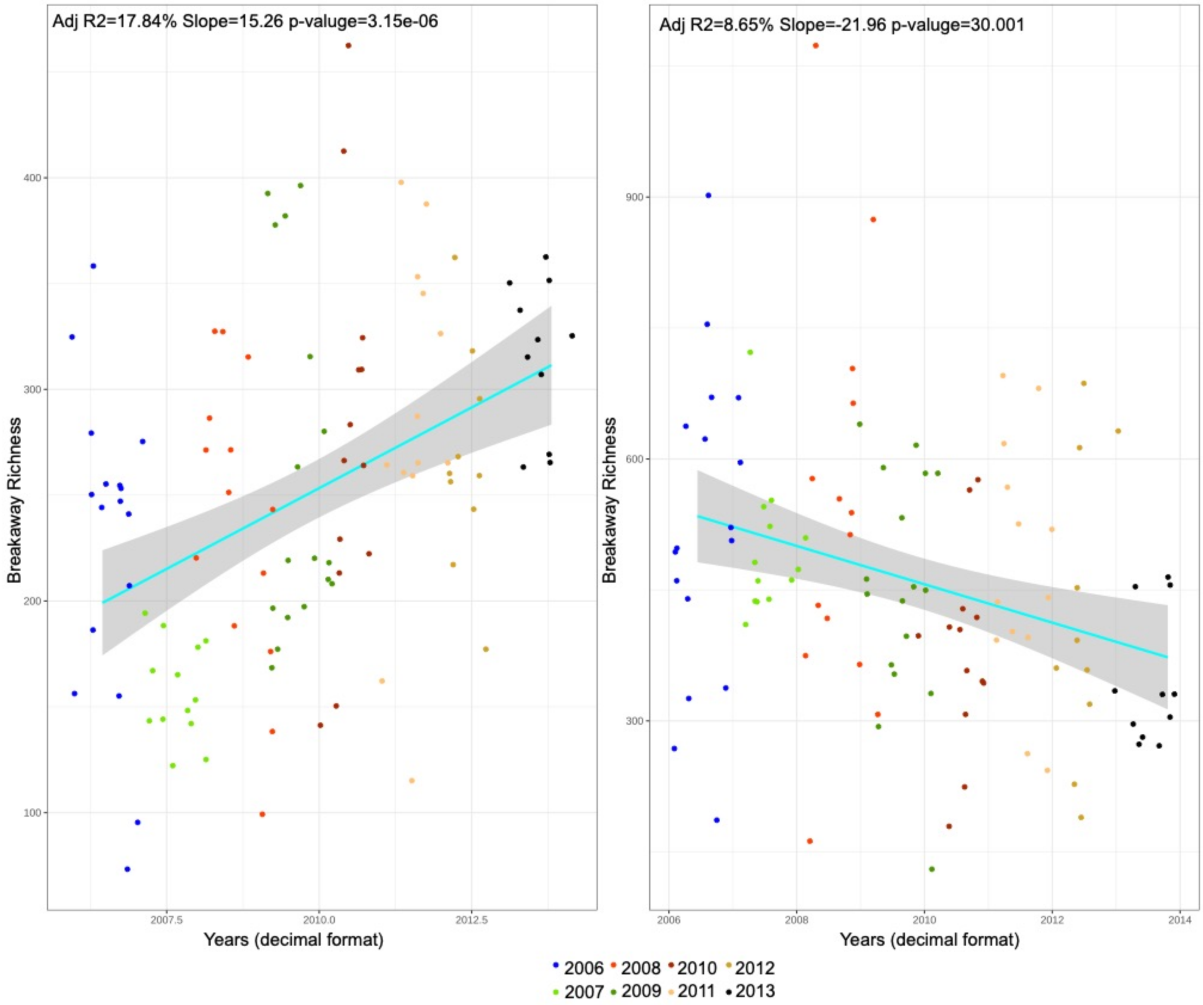


***S5. Breakaway estimate of species richness across years.***

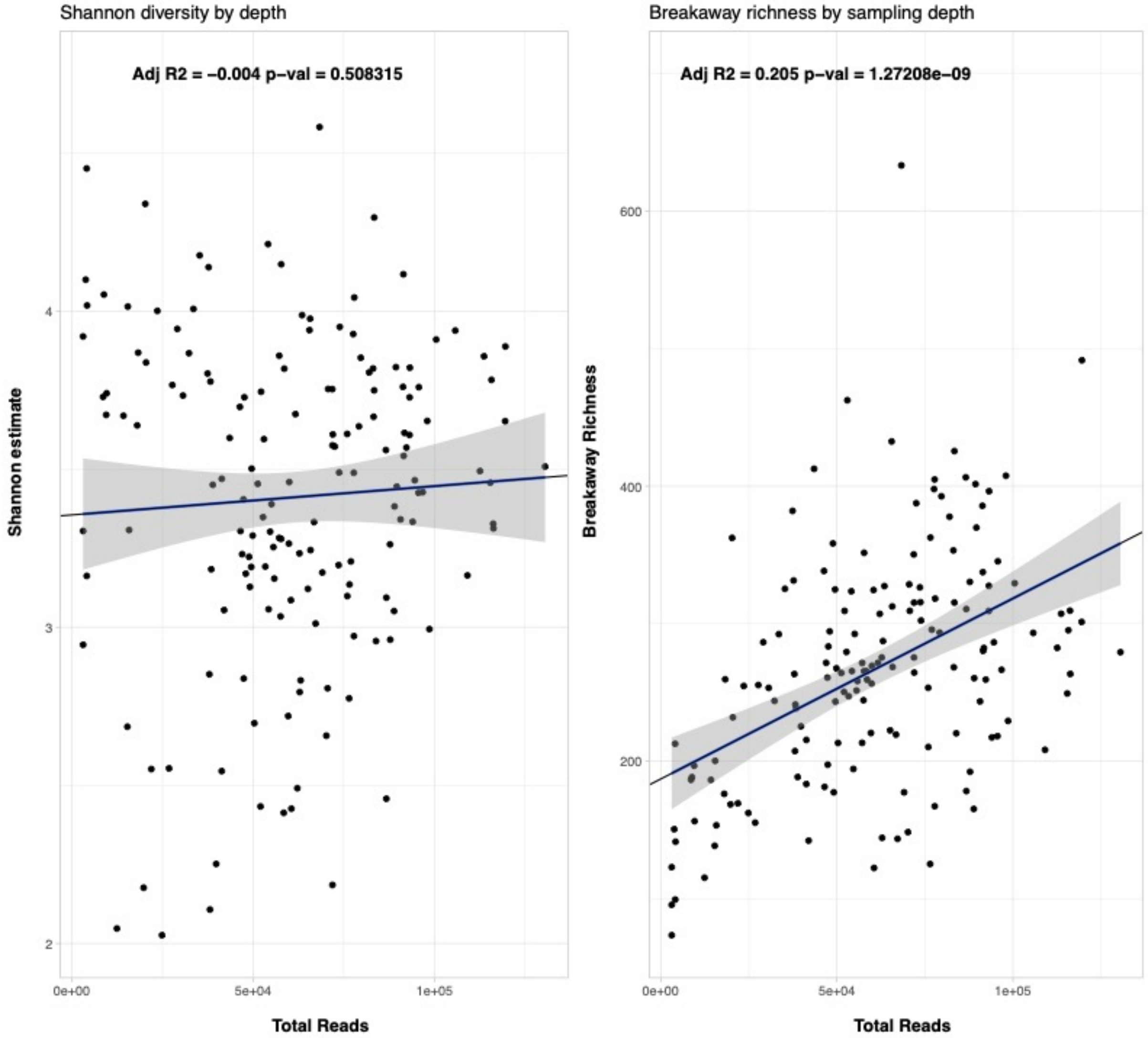


**S6. Diversities by sampling depth**. *A linear regression was performed between Shannon and total richness estimates and sequencing depth (total reads count). The result showed a non-significant relationship with Shannon estimates (left) and a positive correlation with total richness (right).*

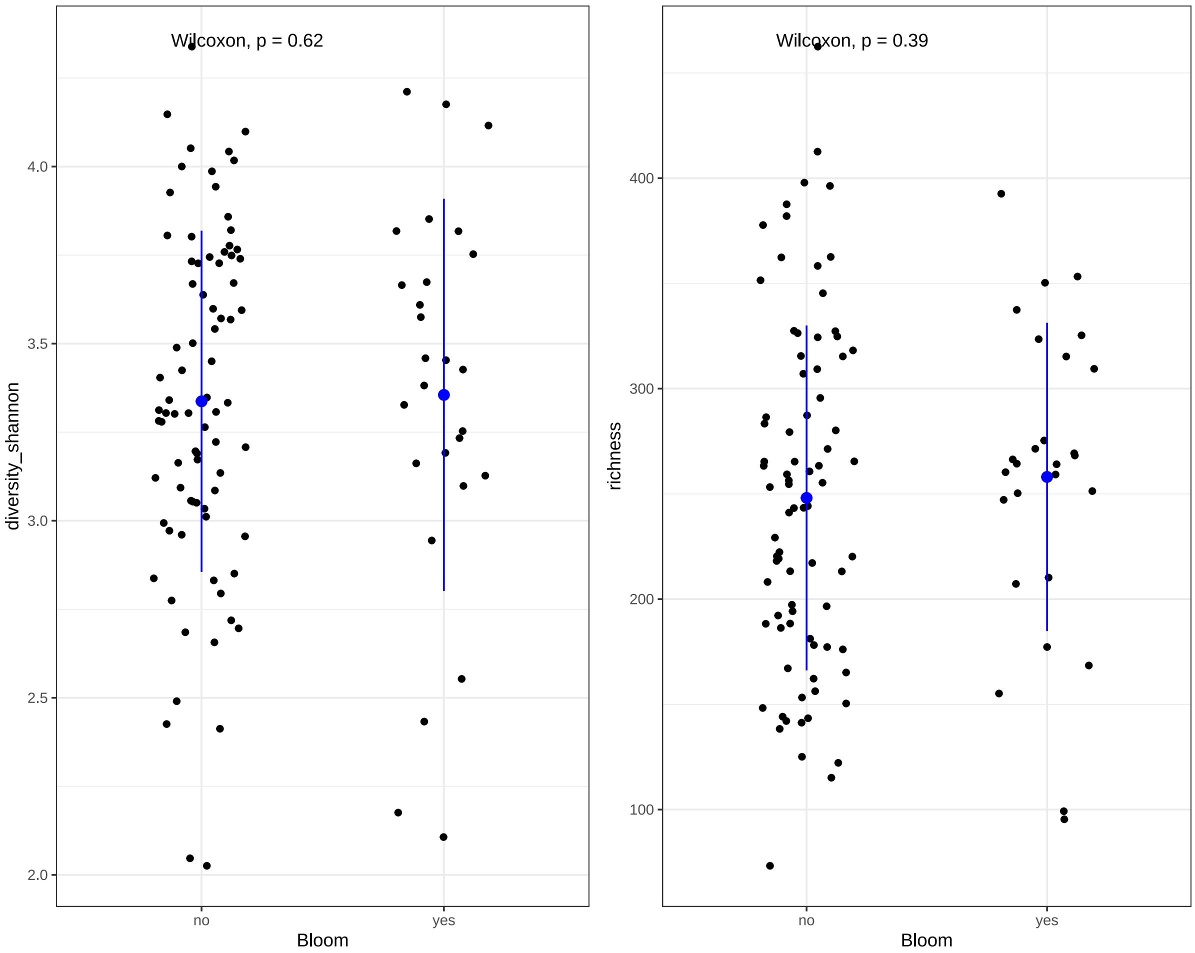


***S7. Diversities estimates during a period of bloom and non-bloom.*** *Non parametric Wilcoxon tests fail to reject the null hypothesis suggesting no significant difference in diversity between bloom and no bloom groups.*

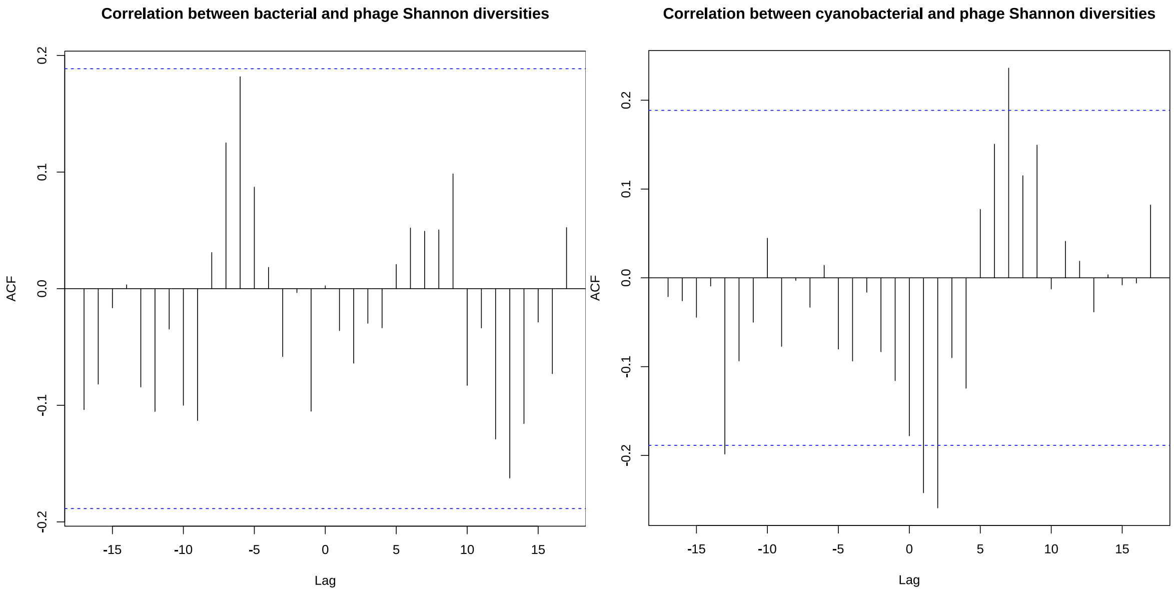

**S8. Cross-correlation analysis between bacterial (left), cyanobacterial (right) and viral Shannon diversity.** Blue lines represent the confidence intervals.

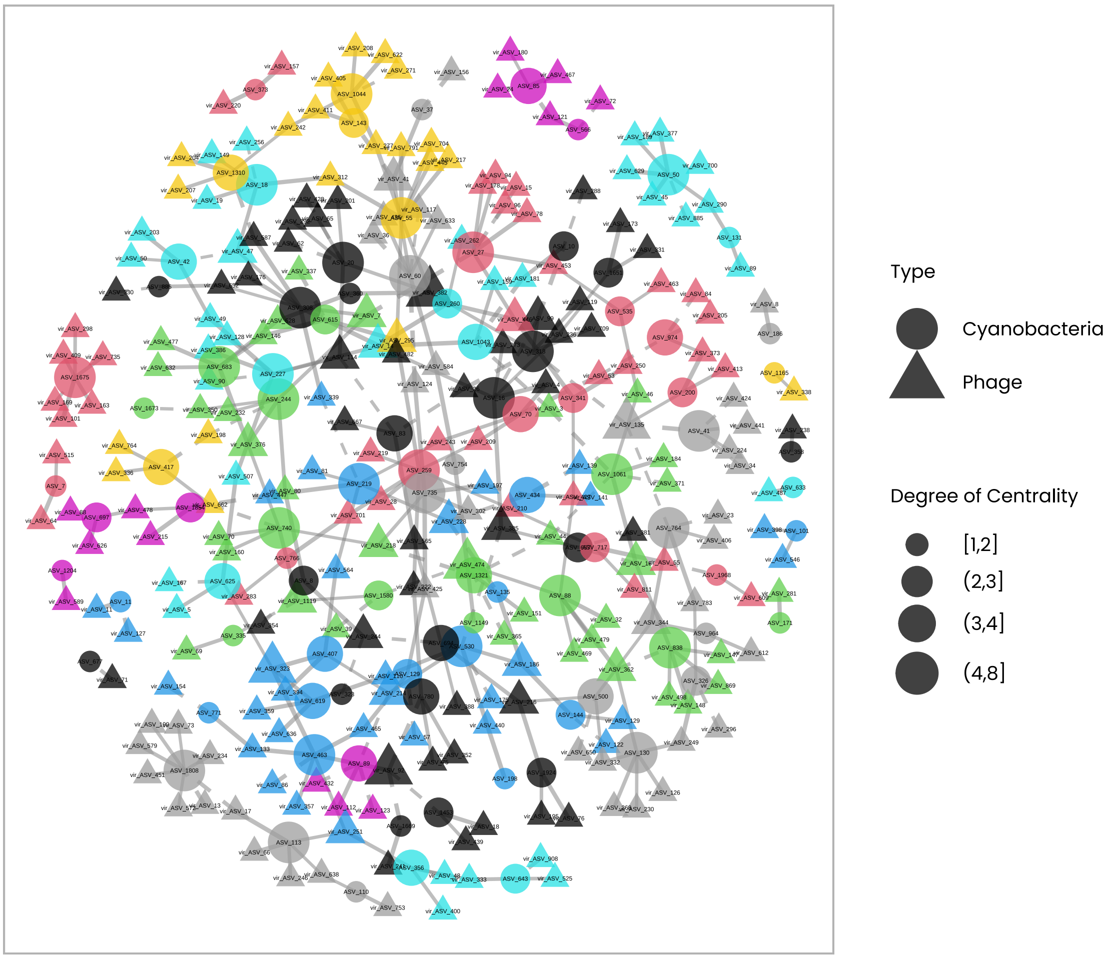


**S9. Phage-Cyanobacteria co-variance network organized by clusters.**

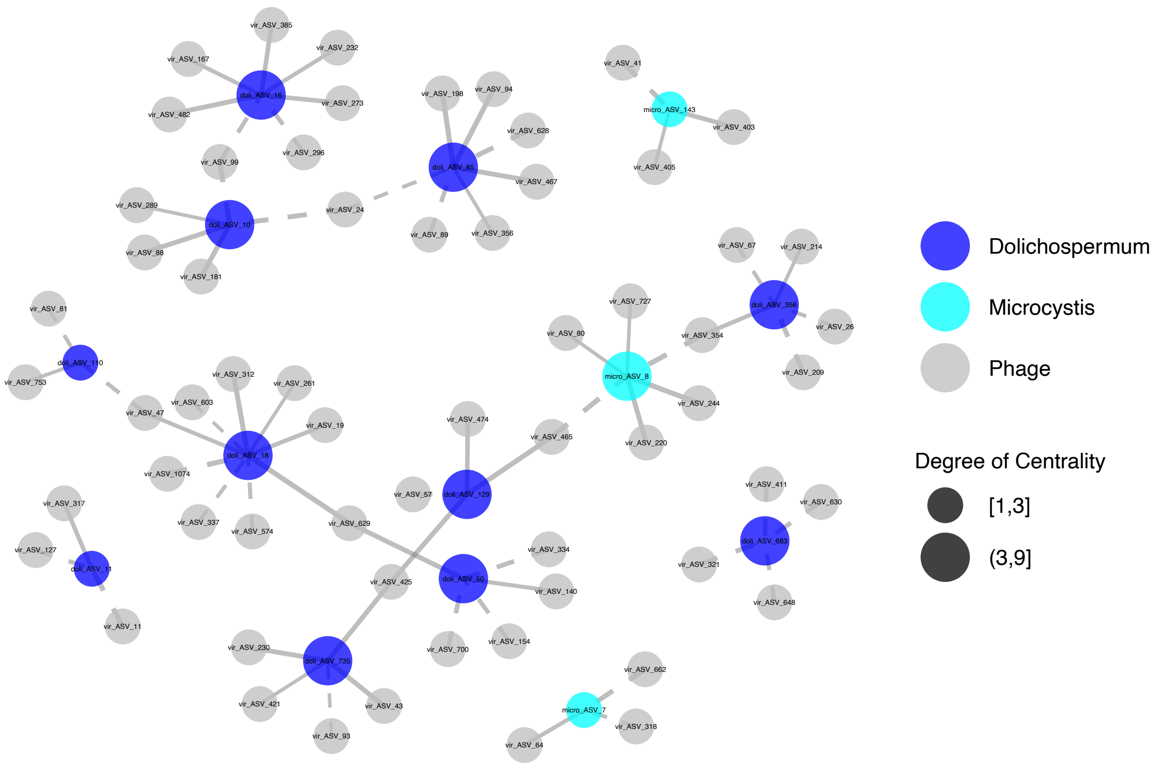


**S10. Phage-*Microcystis*/*Dolichospermum* co-variance network.** Solid and dashed lines correspond to respectively positive and negative co-variance.

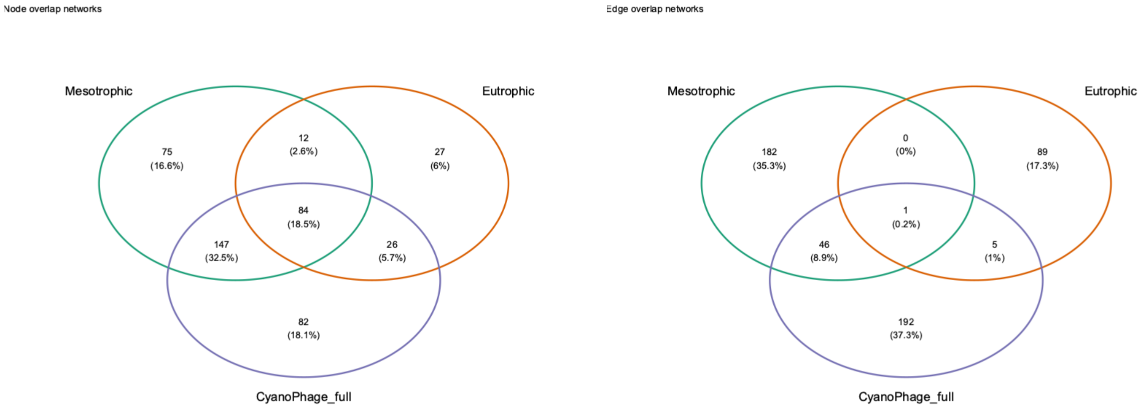


**S11. Comparison of modules (from bipartite network) composition across nutrient conditions (left : nodes; right: edges)**

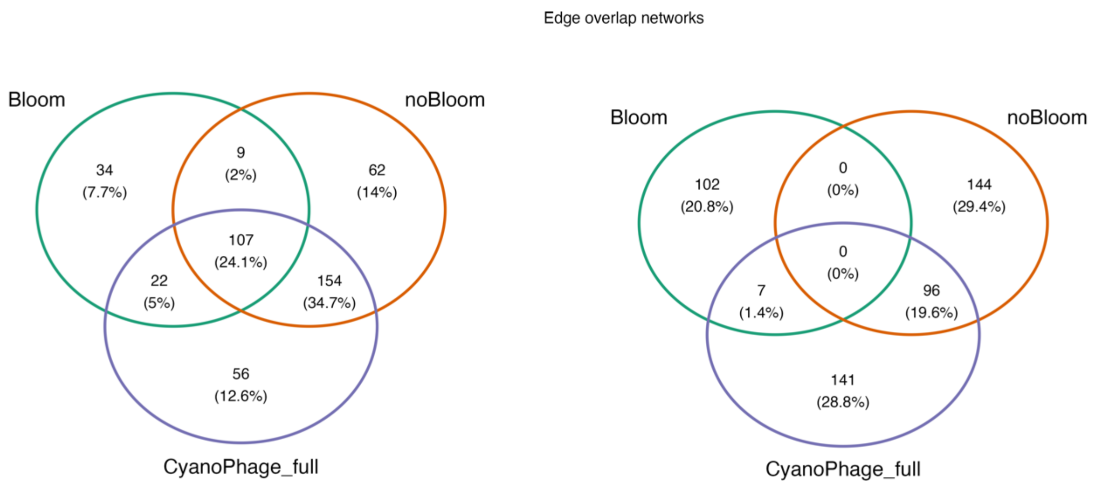


**S12. Comparison of modules (from bipartite network) composition across bloom conditions (left : nodes; right: edges)**

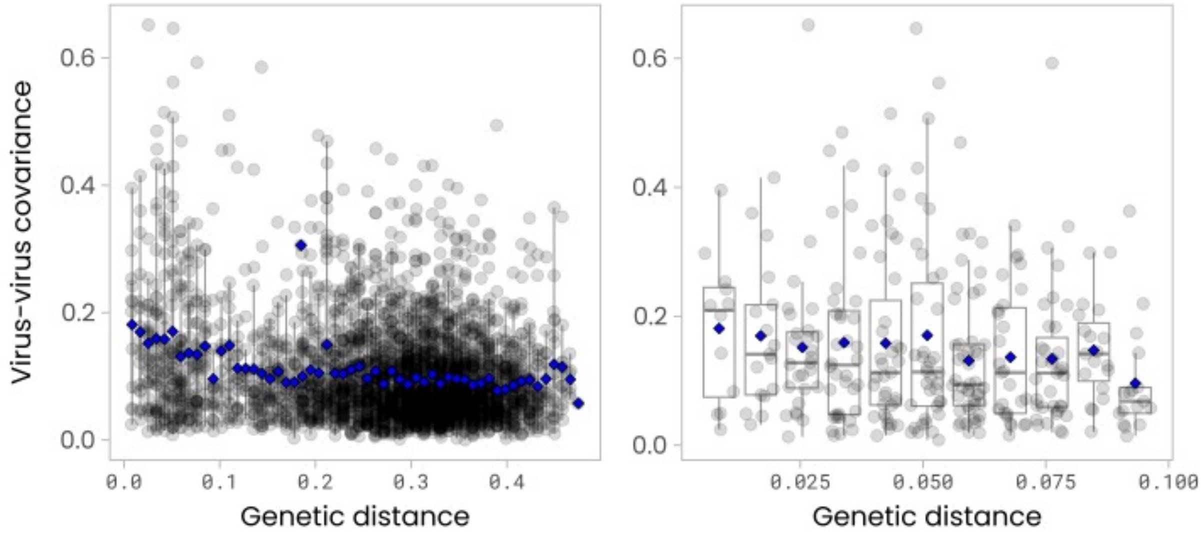


**S13. Viral ASVs co-variance varying across genetic distance.** For each pair of viral ASVs, we compared the value of their genetic distance and co-variance from Spiec-Easi. Linear regression were performed to evaluate the relationship between co-variance and genetic distance: 50-100%: Adjusted R-squared: 0.03857, F-statistic: 122.7 on 1 and 3033 DF, p-value: < 2.2e-16. 90-100%: Adjusted R-squared: 0.0124, F-statistic: 4.866 on 1 and 307 DF, p-value: 0.02813

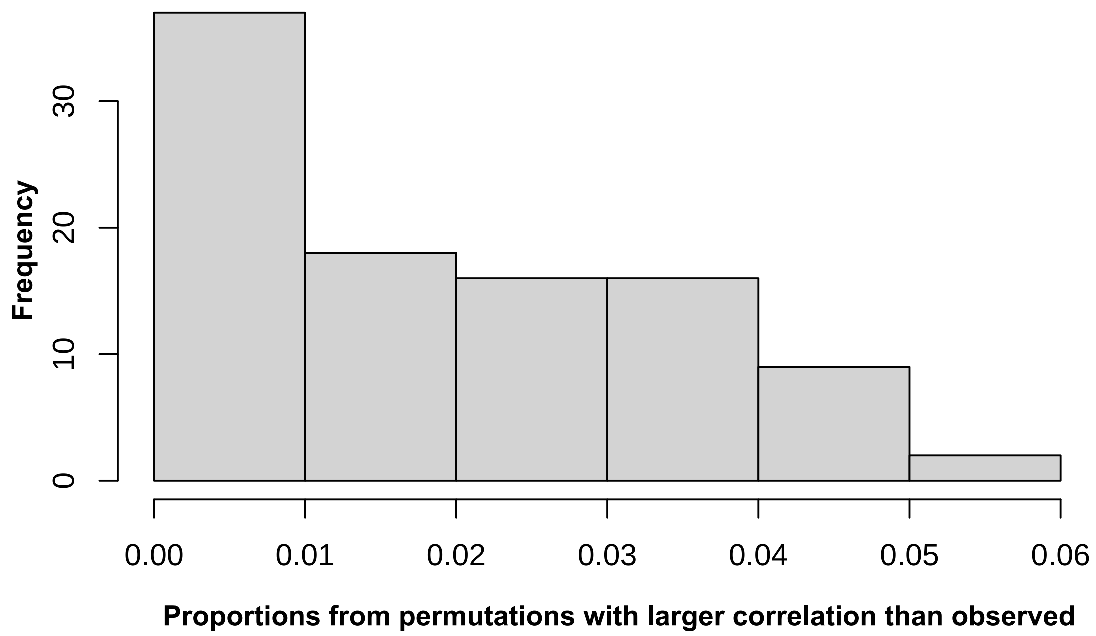


**S14. Permutation test of the association between pair of viral ASVs genetic distance and co-variance with bacterial taxa.** We estimated the proportion (p) of false positive correlations between genetic distance of viral ASVs and the co-variance with bacterial ASVs. The distribution is based on 1000 permutations for each of 98 bacterial ASVs, randomizing the association between |Δr| and genetic distance (Methods). We then calculated p as the number of permutations yielding a larger correlation than observed (numerator) divided by the total number of permutations (denominator).

| VARIABLES | PERMANOVA $R^2$ | PERMANOVA P-VALUE | DISPERSION P-VALUE | PERMANOVA $R^2$ | PERMANOVA P-VALUE | DISPERSION P-VALUE |
|---|---|---|---|---|---|---|
| | Viral ASV | | | Bacterial ASV | | |
| **SITE** | 0.53% | 0.701 | 0.564 | 0.60% | 0.001 | 0.865 |
| **BLOOM** | 2.42% | 0.003 | 0.003 | 10.9% | 0.001 | 0.028 |
| **DOY** | 28.04% | 0.001 | 0.001 | 36.0% | 0.001 | 0.001 |
| **WEEK** | 33.67% | 0.001 | 0.001 | 45.1% | 0.001 | 0.001 |
| **MONTH** | 18.68% | 0.001 | 0.001 | 22.1% | 0.001 | 0.001 |
| **SEASON** | 7.38% | 0.001 | 0.324 | 13.3% | 0.001 | 0.001 |
| **YEAR** | 28.57% | 0.001 | 0.061 | 12.2% | 0.001 | 0.12 |

**Supplementary Table 1. Temporal variables explaining phage and bacterial communities.**

| | MESOTROPHIC | EUTROPHIC | FULL NETWORK |
|---|---|---|---|
| **NODES** | | | |
| **MESOTROPHIC** | 0 | 0.74 | 0.46 |
| **EUTROPHIC** | 0.74 | 0 | 0.71 |
| **FULL NETWORK** | 0.46 | 0.71 | 0 |
| **EDGES** | | | |
| **MESOTROPHIC** | 0 | 0.99 | 0.89 |
| **EUTROPHIC** | 0.99 | 0 | 0.98 |
| **FULL NETWORK** | 0.89 | 0.98 | 0 |

**Supplementary Table 3.** Dissimilarity (Jaccard) between nodes composition across nutrient conditions. A value of 0 indicates an identical composition.

| | BLOOM | NO BLOOM | FULL NETWORK |
|---|---|---|---|
| **NODES** | | | |
| **BLOOM** | 0 | 0.70 | 0.66 |
| **NO BLOOM** | 0.70 | 0 | 0.36 |
| **FULL NETWORK** | 0.66 | 0.36 | 0 |
| **EDGES** | | | |
| **BLOOM** | 0 | 1 | 0.98 |
| **NO BLOOM** | 1 | 0 | 0.75 |
| **FULL NETWORK** | 0.98 | 0.75 | 0 |

**Supplementary Table 4.** Dissimilarity (Jaccard) between nodes composition across bloom conditions.

| NETWORK | PHAGE-BACTERIA | PHAGE-CYANOBACTERIA |
|---|---|---|
| **EDGES NUMBER** | 2628 | 353 |
| **MEAN CONNECTIONS BACTERIA** | 8.14 | 3.72 |
| **MEAN CONNECTIONS PHAGE** | 16.3 | 1.45 |
| **NUMBER OF BACTERIA WITH AT LEAST 1 SIMILAR CO-VARIANT VIRAL ASV** | 1101 | 79 |

**Supplementary Table 5.** Bacteria and Cyanobacteria co-variance networks with phages.
